# Low-abundance αSyn-112 promotes αSyn-140 aggregation in vitro and forms immunoreactive deposits in Parkinson’s disease brain tissue

**DOI:** 10.64898/2026.07.05.736591

**Authors:** Alexander Röntgen, Giuliana Fusco, Jonathan Breiter, Joseph S. Beckwith, Joanne Lachica, Christina E. Toomey, Jaijeet Singh, Oxana Klementieva, Sonia Gandhi, Steven Lee, Alfonso De Simone, Zenon Toprakcioglu, Michele Vendruscolo

**Affiliations:** Yusuf Hamied Department of Chemistry, University of Cambridge, Cambridge CB2 1EW, United Kingdom; Department of Pharmacy, University of Naples Federico II, Naples 80131, Italy; Department of Clinical and Movement Neurosciences, UCL Queen Square Institute of Neurology, University College London, London, United Kingdom; The Francis Crick Institute, London, United Kingdom; Department of Experimental Medical Science, Lund University, 22180 Lund, Sweden

## Abstract

The aggregation of α-synuclein (αSyn) is a molecular hallmark of Parkinson’s disease (PD) and other synucleinopathies. Understanding the molecular mechanisms that determine the aggregation of this protein may thus facilitate the development of disease-modifying therapies. While αSyn is most commonly expressed as a 140-residue protein (αSyn-140), recent evidence suggests an involvement of alternatively spliced αSyn isoforms in disease onset and progression. Here, we report and characterise the interaction between αSyn-140 and the aggregation-prone αSyn-112 variant, one of the most abundant αSyn splice isoforms. We found that amounts as low as 1% of αSyn-112 accelerate the nucleation and aggregation of αSyn-140. To further investigate this phenomenon, we employed MALDI-MS and NMR spectroscopy, confirming that αSyn-140 and αSyn-112 monomers interact strongly with one another. Furthermore, to assess the association of αSyn-112 with disease pathology, we performed immunohistochemical staining combined with confocal microscopy on PD brain samples. Thereby, we found an increase in the number as well as the area of αSyn-112 immunoreactive aggregates compared to healthy controls. These results illustrate how low-abundance αSyn splice isoforms can modulate the aggregation landscape of αSyn-140 and in turn contribute to the molecular heterogeneity of synucleinopathies.

## Introduction

The pathogenesis of Parkinson’s disease (PD) and other synucleinopathies is linked to the aggregation of α-synuclein (αSyn) from soluble monomers into amyloid fibrils, which are the major components of Lewy pathology in patient brains (*1–4*). αSyn is encoded by the *SNCA* gene and is commonly described as a 140-residue protein (αSyn-140), which is highly abundant in presynaptic nerve terminals and plays a role in neurotransmission (*5, 6*).

The canonical amino acid sequence of αSyn can be divided into three functional regions: an amphipathic N-terminus (residues 1-60), which is important for binding of lipid membranes (*7, 8*), a central hydrophobic non-amyloid β component (NAC) region (residues 61-95), which drives amyloid aggregation (*9–11*), and a highly negatively charged, flexible C-terminus (residues 96-140), which inhibits aggregation (*12, 13*).

Increasing evidence indicates that αSyn does not exist as a single molecular species, but is rather present as a spectrum of proteoforms, including post-translational modifications, truncations and splice variants, with divergent biophysical properties (*14–17*). Already shortly after the discovery of the 140-amino acid αSyn variant (*18*), an additional 112-residue isoform of αSyn (αSyn-112) was identified which lacks the C-terminal exon 5 (residues 103-130) due to alternative splicing of the *SNCA* mRNA (*5*). Subsequent studies have shown that, by exon skipping, a range of further splice isoforms can be generated besides αSyn-112, including αSyn-126 lacking exon 3 (residues 41-54), and αSyn-98 lacking both exons 3 and 5 (*19–22*). Importantly, αSyn-112 was recently found to be one of the most strongly transcribed splice isoforms after αSyn-140, and was detected at the protein level in human brain tissue and blood using long-read transcriptomics and top-down proteomics, respectively (*19, 23, 24*). Due to its lack of the C-terminal exon 5 sequence, αSyn-112 exhibits enhanced aggregation kinetics compared to αSyn-140, and forms morphologically distinct, clump-like aggregates consisting of amyloid fibrils *in vitro* (*25*). Moreover, previous work has shown that αSyn-112 can interact with αSyn-140 and may therefore act as a pathological factor in the development of synucleinopathies (*25*).

To obtain a more accurate understanding of the interplay between αSyn-112 and αSyn-140, and to assess the disease relevance of alternatively spliced αSyn, we therefore performed a detailed characterisation of the molecular interaction and co-aggregation between αSyn-112 and αSyn-140. Using thioflavin T (ThT) aggregation assays combined with chemical kinetics and transmission electron microscopy (TEM), we assessed the microscopic steps in the aggregation network that are affected by the addition of small amounts of αSyn-112. We then used matrix-assisted laser desorption/ionisation mass spectrometry (MALDI-MS), solution-state nuclear magnetic resonance (NMR) spectroscopy and optical photothermal infrared (OPTIR) spectroscopy to gain insights into the interaction of the two isoforms at the molecular level. To investigate the possible role of αSyn-112 in disease, we performed immunohistochemical (IHC) staining of human brain tissue, showing that αSyn-112-immunoreactive aggregates are more prominent in PD compared to healthy brain tissue. Overall, our study emphasises the importance to further understand the effects of alternative splicing in neurodegeneration if we aim to develop effective therapies that target the disease mechanism.

## Results

### αSyn-112 enhances the aggregation of αSyn-140

We first sought to explore how αSyn-112 affects the aggregation of αSyn-140 and elucidate the microscopic kinetic rates. We incubated the pure αSyn-140 system (100% αSyn-140) and the mixed αSyn-140+αSyn-112 system (90% αSyn-140 + 10% αSyn-112) at varying total αSyn concentrations (5-25 µM), and monitored the aggregation over time by adding the amyloid-binding dye thioflavin T (ThT). Thereby, we determined the kinetic rate constants of the aggregation reaction (**Figure 1**). This experiment revealed that the mixed system undergoes aggregation on shorter timescales than the pure system at all total αSyn concentrations tested (**Figure 1A-D**). Furthermore, based on the aggregation half-times (t_1/2_), we determined the scaling exponents for the pure and mixed aggregation systems. In both cases, the values we determined were consistent with a *Saturating Elongation and Fragmentation* model of aggregation (**Figure 1E, F**) (*25, 26*). Utilising the platform AmyloFit, we fitted the aggregation data accordingly and derived the kinetic rate constants of the aggregation process for each system. Our analysis revealed a marked elevation in the combined rate constants: the value for k_+_k_n_ (*i.e.* primary nucleation and elongation) was increased by a factor of 4.1, while the value for k_+_k_-_ (*i.e.* elongation and fragmentation) was increased by a factor of 10.4, upon addition of αSyn-112. These results reveal that low amounts of αSyn-112 can strongly enhance the aggregation of αSyn-140 through an increase in the kinetic rate constants.

**Figure 1.**
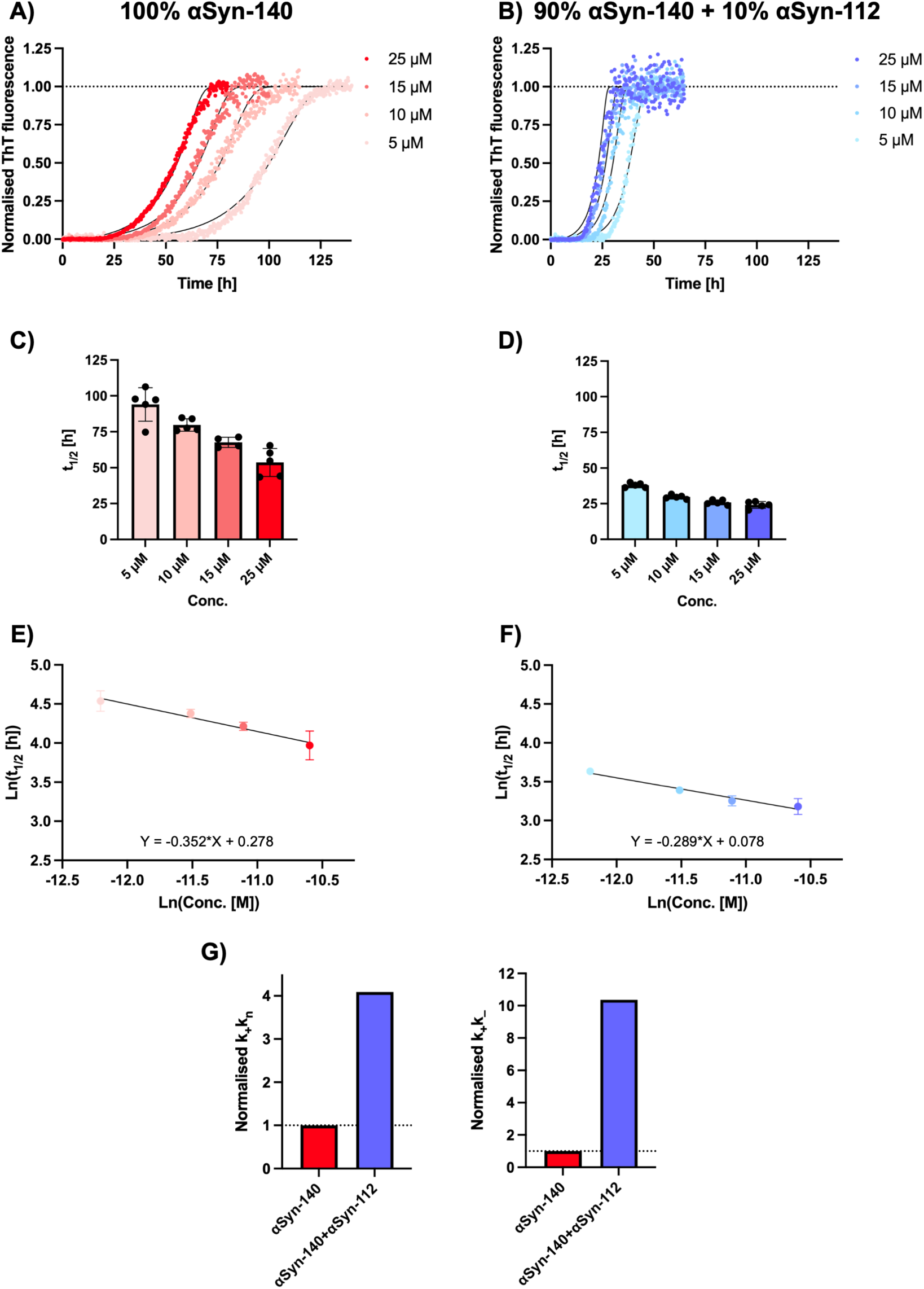
Low amounts of αSyn-112 increase the microscopic rate constants of αSyn-140 aggregation. **(A,B)** The aggregation of 100% αSyn-140 (A) and 90% αSyn-140 + 10% αSyn-112 (B) at varying total protein concentrations was assessed by monitoring ThT fluorescence over time. Normalised traces were fitted to the *Saturating Elongation and Fragmentation* model (solid lines) using AmyloFit. **(C, D)** The aggregation half-times (t_1/2_) exhibit a characteristic dependence on protein concentration. **(E, F)** Using double-logarithmic plots, the scaling exponents, *i.e.* the slopes of the linear regression functions, were determined. Data are shown as mean ± SD. **(G)** Kinetic parameters derived from the fitting of the aggregation data in (A, B): k_+_k_n_ (primary nucleation and elongation), k_+_k_–_(elongation and fragmentation).

We next investigated the effect of varying the amount of αSyn-112 on the aggregation of αSyn-140. By incrementally decreasing the percentage of αSyn-112, we were able to identify the lowest amount of αSyn-112 required to elicit a significant acceleration in the aggregation process. As αSyn splice isoforms are estimated to account for only a small proportion of the overall αSyn protein content in cells (*19*), we performed co-aggregation reactions of mixtures containing 0%-10% αSyn-112 with 100%-90% αSyn-140, respectively. The total αSyn concentration was kept constant while varying the percentage of αSyn-112. The experiments were performed at both 25 and 5 µM total αSyn, *i.e.* at the highest and lowest concentrations tested in the previous experiment (**Figure 1**). This was done to identify potential changes in the effect of αSyn-112 at different total protein concentrations. Moreover, these concentrations are in line with studies estimating the total αSyn concentration at synapses to be in the low to intermediate micromolar range (*27, 28*).

We found that increasing amounts of αSyn-112 gradually accelerated the aggregation of αSyn-140 (**Figure 2A,B**). To quantify the change in the aggregation rate, we first determined the aggregation half-times (t_1/2_). We found that, at 25 µM total αSyn, the t_1/2_ value was reduced by 35%, from 37 h (0% αSyn-112) to 24 h (10% αSyn-112). Interestingly, at 5 µM total αSyn concentration, the accelerating effect relative to the control condition was even stronger, where we observed a reduction in t_1/2_ by 66%, from 94 h (0% αSyn-112) to 32 h (10% αSyn-112). We should note that 1% αSyn-112 was sufficient to elicit a statistically significant reduction in t_1/2_ at 5 µM total αSyn, whereas 5% αSyn-112 was required to achieve a significant reduction in t_1/2_ at 25 µM total αSyn. This indicates the speed of aggregation scales more strongly with the αSyn-112 ratio at lower total αSyn concentrations. Thereby, despite a 5-fold difference in the total protein concentration (5 µM and 25 µM), the aggregation half-times converge to similar values with increasing percentages of αSyn-112. This could be because the aggregation process may be less saturated at lower total protein concentrations, whereby adding an aggregation-promoting isoform has a relatively larger effect on the system.

**Figure 2.**
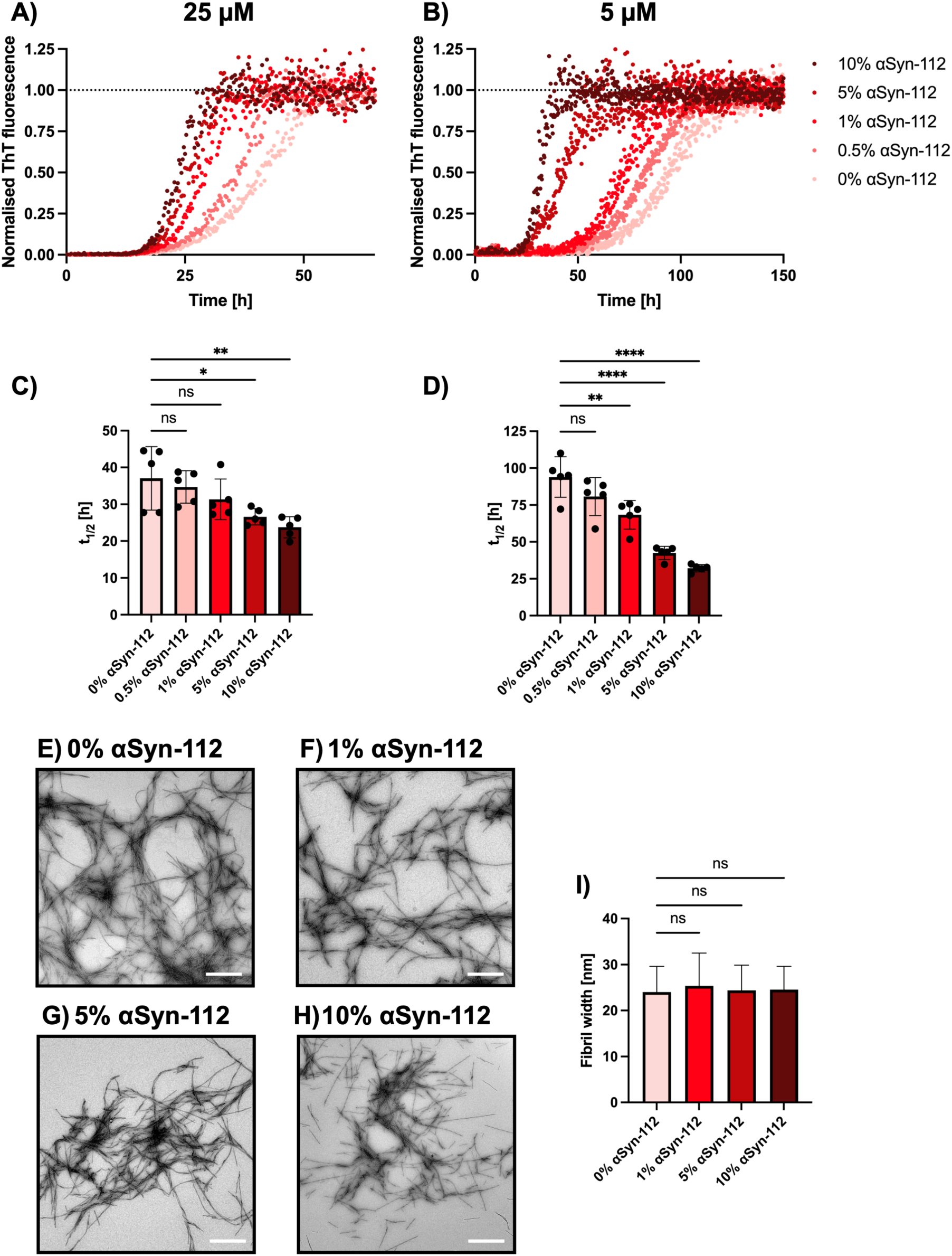
As little as 1% αSyn-112 accelerates αSyn-140 aggregation without altering fibril morphology. **(A,B)** The co-aggregation kinetics of mixtures containing 100–90% αSyn-140 with 0–10% αSyn-112, respectively, at 25 µM (A) and 5 µM (B) total αSyn concentrations was assessed by monitoring ThT fluorescence. **(C,D)** Half-time plot as a function of αSyn-112 concentration for 25 µM total αSyn concentration (C) and 5 µM (D) total αSyn concentration. Data are shown as mean ± SD. **(E-H)** Representative images of amyloid fibrils formed by αSyn-140 co-aggregated with αSyn-112. Scale bars = 1 µm. **(I)** The fibrils of all conditions tested displayed similar widths. **(C,D,I)** One-way ANOVA with Dunnett’s post-hoc test. ****p<0.0001, **p<0.01, *p<0.05, ns = non-significant.

We further visualised the resulting aggregates using transmission electron microscopy (TEM) to assess the aggregate morphology. We found that elongated amyloid fibrils with similar widths were formed at all ratios of αSyn-112 tested (**Figure 2E-I**). This is in contrast to the large fibril clumps observed for pure αSyn-112, which are several micrometres in diameter (*25*). Given that no such clumps were observed in the co-aggregation reactions, these data support the idea that αSyn-112 monomers were incorporated into the amyloid fibrils along with αSyn-140.

### αSyn-140 and αSyn-112 form co-oligomers

Following our finding that the accelerated aggregation between αSyn-140 and αSyn-112 may be linked to an increase in primary nucleation, we asked whether the two isoforms form co-oligomers during the aggregation process. To address this question, we used matrix-assisted laser desorption/ionisation mass spectrometry (MALDI-MS), which is sensitive to low-mass assemblies, to analyse the oligomeric species formed in pure αSyn-140 (100%), pure αSyn-112 (100%) and mixed αSyn-140 + αSyn-112 (90% + 10%) reactions after 20 h of incubation (**Figure 3**). As expected, in the reactions containing pure αSyn-140, or pure αSyn-112, we exclusively observed the formation of the respective homo-oligomers (**Figure 3A,B**). However, in the mixed reaction, we detected αSyn-140+αSyn-112 hetero-dimers alongside αSyn-140 homo-oligomers (**Figure 3C**). Importantly, no αSyn-112 homo-dimers were observed in the mixed sample. This clearly suggests that, when incubated with an excess of αSyn-140, αSyn-112 preferentially binds to αSyn-140 and is therefore more likely to form hetero-dimers with αSyn-140 as opposed to αSyn-112 homo-dimers. The lower intensity of the hetero-dimer signal compared with αSyn-140 homo-oligomers is consistent with the low proportion of αSyn-112 in the reaction. Together, these results provide evidence that the two αSyn isoforms co-assemble into hetero-oligomers during the early phase of aggregation and further support our hypothesis that enhanced molecular interactions and heterotypic primary nucleation contribute to the accelerated aggregation of αSyn-140 with αSyn-112.

**Figure 3.**
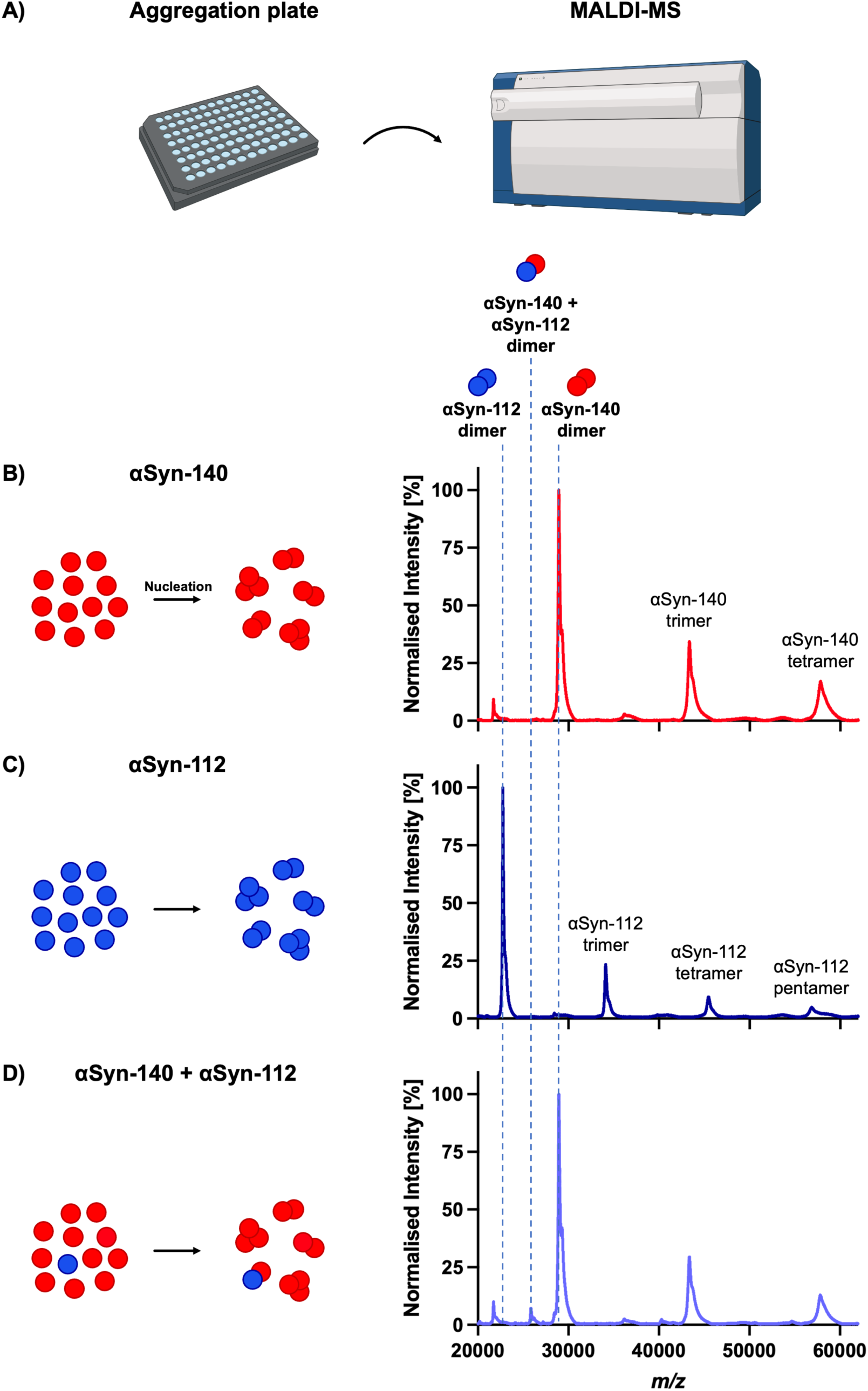
αSyn-140 and αSyn-112 form hetero-oligomers during the early phase of aggregation. **(A)** Schematic workflow. **(B-D)** Aggregation reactions of 100% αSyn-140 (B), αSyn-112 (C), and 90% αSyn-140 with 10% αSyn-112 (D). All samples were analysed using MALDI-MS following a 20 h incubation. Parts of this figure were adapted using BioRender.com.

### NMR reveals that monomer-monomer interactions between αSyn-140 and αSyn-112 are enhanced in the NAC region

To unravel the molecular mechanism that underlies the marked acceleration of aggregation, we investigated the interaction between monomeric αSyn-140 and αSyn-112 using nuclear magnetic resonance (NMR) spectroscopy. We hypothesised that the deletion of most of the negatively charged C-terminus in αSyn-112 promotes the interaction between αSyn-112 and αSyn-140 monomers. Consequently, the closer proximity between the molecules would lead to enhanced primary nucleation and faster amyloid aggregation.

To probe this interaction and to identify the involved regions in both isoforms, we created two pairs of protein constructs. We placed MTSL spin labels in either the NAC region (position 90) or the N-terminus (position 24) of both ^14^N-αSyn-112 and ^14^N-αSyn-140 (**Figure 4A,B**). Next, we individually mixed each of these four constructs with ^15^N-isotope-labelled αSyn-140 (^15^N-αSyn-140) and performed interchain paramagnetic relaxation enhancement (PRE) experiments. Using this method, we determined the relaxation time, T_2_, of the residues in the ^15^N-αSyn-140 molecule in the presence of any of the four MTSL-labelled ^14^N-αSyn species. Thereby, we can deduce which residues in the ^15^N-αSyn-140 interact more strongly with the label-bearing region in the ^14^N-αSyn-MTSL construct. This approach therefore allows us to compare the interactions of the ^15^N-αSyn-140 molecule with the ^14^N-αSyn constructs.

**Figure 4.**
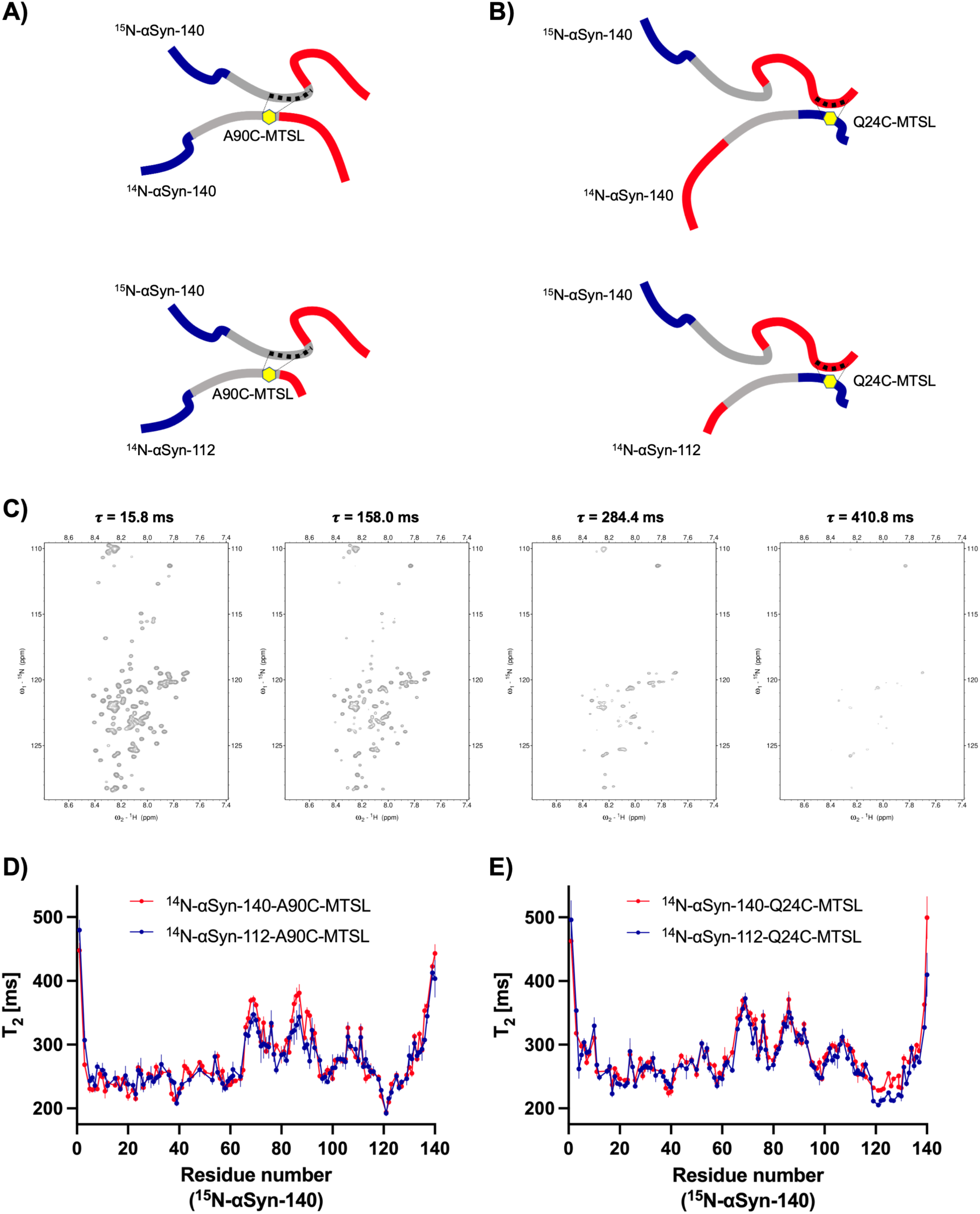
Monomer–monomer interactions between αSyn-140 and αSyn-112 are enhanced in the NAC region. **(A,B)** Schematic representation of the mixtures compared in this experiment. **(C)** Representative NMR spectra of ^15^N-αSyn-140 mixed with ^14^N-αSyn-112-Q24C-MTSL using different relaxation delays, τ, to determine the relaxation times, T_2_, of the amino acid residues in ^15^N-αSyn-140. **(D,E)** Relaxation times, T_2_, for the individual amino acid residues in ^15^N-αSyn-140 when mixed with (D) ^14^N-αSyn-140-A90C-MTSL (red), ^14^N-αSyn-112-A90C-MTSL (blue), (E) ^14^N-αSyn-140-Q24C-MTSL (red) or ^14^N-αSyn-112-Q24C-MTSL (blue). Data are shown as mean ± SD.

Representative NMR spectra, acquired using different relaxation delays, τ, are shown in **Figure 4C**. Using this strategy, we found that residues located in the NAC region of ^15^N-αSyn-140 interact more strongly with the NAC region of αSyn-112 (^14^N-αSyn-112-A90C-MTSL) than with the NAC region of αSyn-140 (^14^N-αSyn-140-A90C-MTSL), which is indicated by lower T_2_ values of the involved residues in ^15^N-αSyn-140 (**Figure 4D**). This finding supports our hypothesis from the above kinetic experiments that, at the monomeric level, αSyn-140 interacts more strongly with αSyn-112 than with other αSyn-140 molecules. Moreover, the results are in line with the fact that the hydrophobic NAC region is well-known to be a key driver of αSyn amyloid aggregation (*9–11*). In addition, this series of experiments revealed that the C-terminus of ^15^N-αSyn-140 interacts more strongly with the N-terminus of αSyn-112 (^14^N-αSyn-112-Q24C-MTSL) than with that of αSyn-140 (^14^N-αSyn-140-Q24C-MTSL) (**Figure 4E**). Notably, the decrease in T_2_ was observed for residues located in the C-terminal region of ^15^N-αSyn-140 (from residue 119 onwards), indicating that the αSyn-140 population interacts more favourably in a head-to-tail orientation with αSyn-112 than with αSyn-140.

### Hybrid αSyn-140+αSyn-112 fibrils are structurally distinct but possess similar seeding potency compared to pure αSyn-140 fibrils

We next assessed whether the interaction between αSyn-140 and αSyn-112 results in differences in the secondary structure of the amyloid fibrils upon co-aggregation. To this end, we compared pure αSyn-140 fibrils, pure αSyn-112 fibrils, and hybrid αSyn-140+αSyn-112 fibrils, using optical photothermal infrared (OPTIR) spectroscopy to detect structural differences (**Figure 5A**), establishing structural fingerprints displayed as a heatmap in **Figure 5B**. While all three types of aggregates possess profiles characteristic of amyloid fibrils (*i.e.* displaying a shift in the Amide I peak from ∼1650 cm^−1^ to ∼1630 cm^−1^), we found that hybrid αSyn-140+αSyn-112 fibrils exhibit several features which more closely resemble pure αSyn-140 (*i.e.* similar peaks at 1545, 1656, 1676 and 1692 cm^−1^), rather than αSyn-112 fibrils. However, despite sharing several features with pure aggregates, the hybrid αSyn-140+αSyn-112 aggregates also possess distinct structural characteristics (*i.e.* low peak area at 1664 cm^−1^). Consequently, these hybrid fibrils have an overall unique secondary structure as a result of the co-aggregation of the two isoforms.

**Figure 5.**
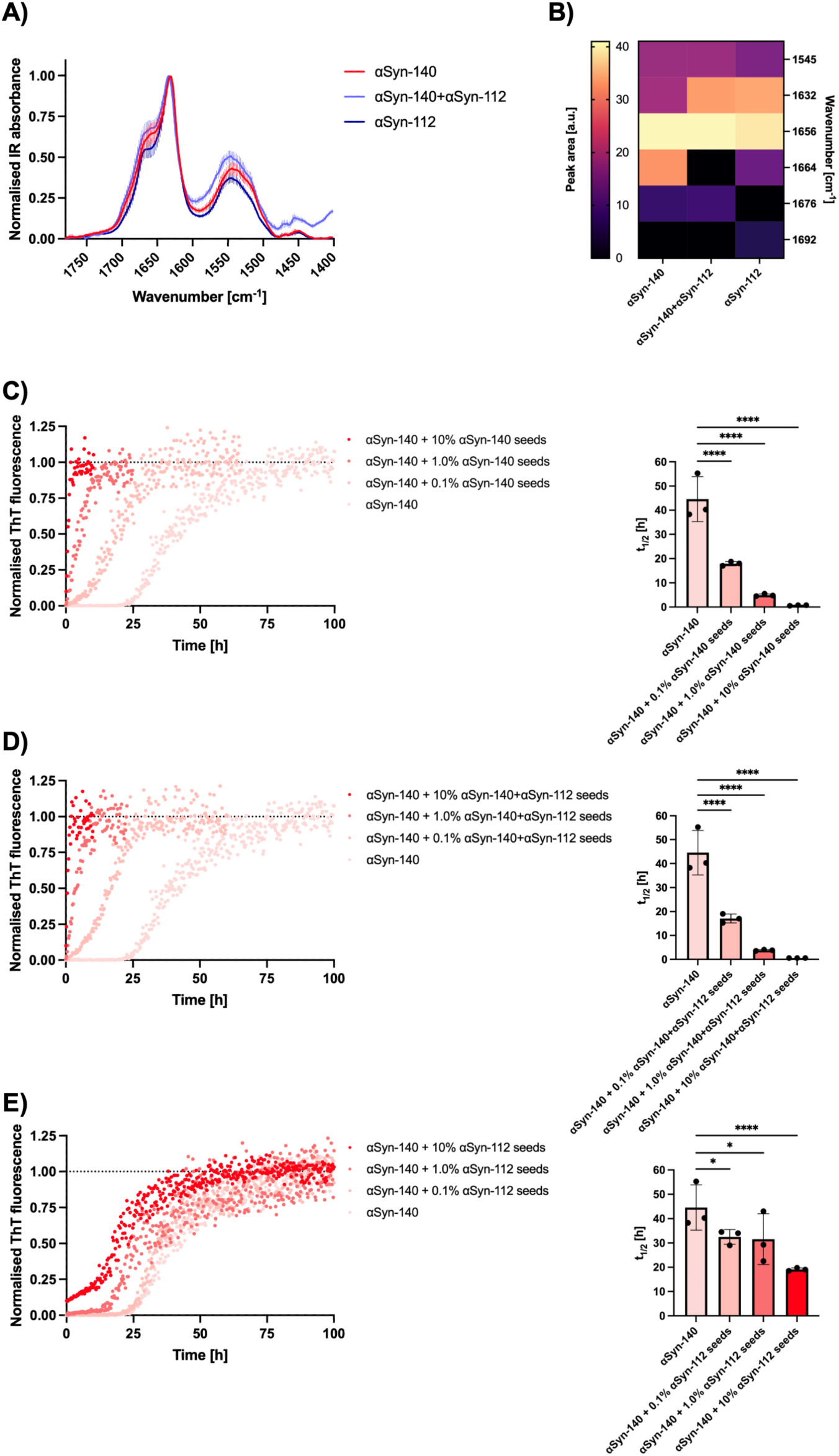
Hybrid αSyn-140+αSyn-112 fibrils are structurally distinct but seed αSyn-140 as potently as pure αSyn-140 fibrils. **(A)** Averaged and normalised OPTIR spectra of αSyn fibrils. **(B)** Heatmap displaying structural fingerprints based on the area under the peaks. **(C-E)** Aggregation traces of αSyn-140 monomers mixed with pre-formed fibrils seeds composed of (C) 100% αSyn-140, (D) 90% αSyn-140 and 10% αSyn-112, and (E) 100% αSyn-112 (**left panels**) and the corresponding t_1/2_ values (**right panels**). Data are shown as mean ± SD. One-way ANOVA with Dunnett’s post-hoc test. ****p<0.0001, *p<0.05.

In recent years, the spread of αSyn fibril seeds between cells has gained increasing attention as a mechanism by which αSyn pathology propagates through the brain (*29*). We therefore investigated how potent hybrid αSyn-140+αSyn-112 fibrils would be in kinetic experiments as fibril seeds for αSyn-140 monomers. Again, we compared the hybrid fibrils with pure fibrils of both αSyn-140 and αSyn-112. For this, αSyn-140 monomers were incubated with 0%, 0.1%, 1.0% or 10% of seeds, consisting either of 100% αSyn-140 (**Figure 5C**), 90% αSyn-140 + 10% αSyn-112 (**Figure 5D**), or 100% αSyn-112 (**Figure 5E**). The results demonstrate that pure αSyn-140 and hybrid αSyn-140+αSyn-112 fibrils are equally potent at seeding the aggregation of αSyn-140 monomers, in both cases leading to a marked reduction of t_1/2_ by 60% (0.1% seeds), 90% (1.0% seeds) and 99% (10% seeds) (**Figure 5C,D**). In contrast, the effect was much diminished when incubating αSyn-140 monomers with pure αSyn-112 fibrils, resulting merely in a reduction of t_1/2_ by 27% (0.1% seeds), 29% (1.0% seeds) and 57% (10% seeds) (**Figure 5E**). In line with this, it has been previously shown that seeding of αSyn aggregation is most efficient when monomers are mixed with fibril seeds of the same variant (*13, 25*). Notably, we observe identically potent seeding capacities of αSyn-140 and hybrid αSyn-140+αSyn-112 fibrils despite differences in composition and secondary structures. Conversely, αSyn-112 fibril seeds are significantly less compatible with αSyn-140 monomers, leading to a decreased seeding capacity.

### αSyn-112-immunoreactive aggregates are enriched in PD brain tissue

Our *in vitro* experiments demonstrated that small amounts of αSyn-112 accelerate the aggregation of αSyn-140, that the two isoforms form hetero-oligomers and interact through the NAC region, and that the resulting hybrid fibrils seed αSyn-140 monomers as efficiently as pure αSyn-140 fibrils. We therefore asked whether αSyn-112 aggregates are present in diseased tissue, reasoning that if αSyn-112 contributes to αSyn aggregation *in vivo*, it might itself be deposited in patient brains. To test this possibility, we performed immunohistochemical (IHC) staining for αSyn-112 on post-mortem cingulate cortex from four PD patients (Braak stage 6) and four healthy controls, and then imaged the sections using spinning-disk confocal microscopy with automated single-aggregate detection (**Figure 6A**)(*30*). We chose samples from the cingulate cortex as this brain region is known to be heavily affected by Lewy pathology at Braak stage 6. Sections were co-stained for αSyn-140 as a reference for the established pattern of αSyn-140 pathology.

**Figure 6.**
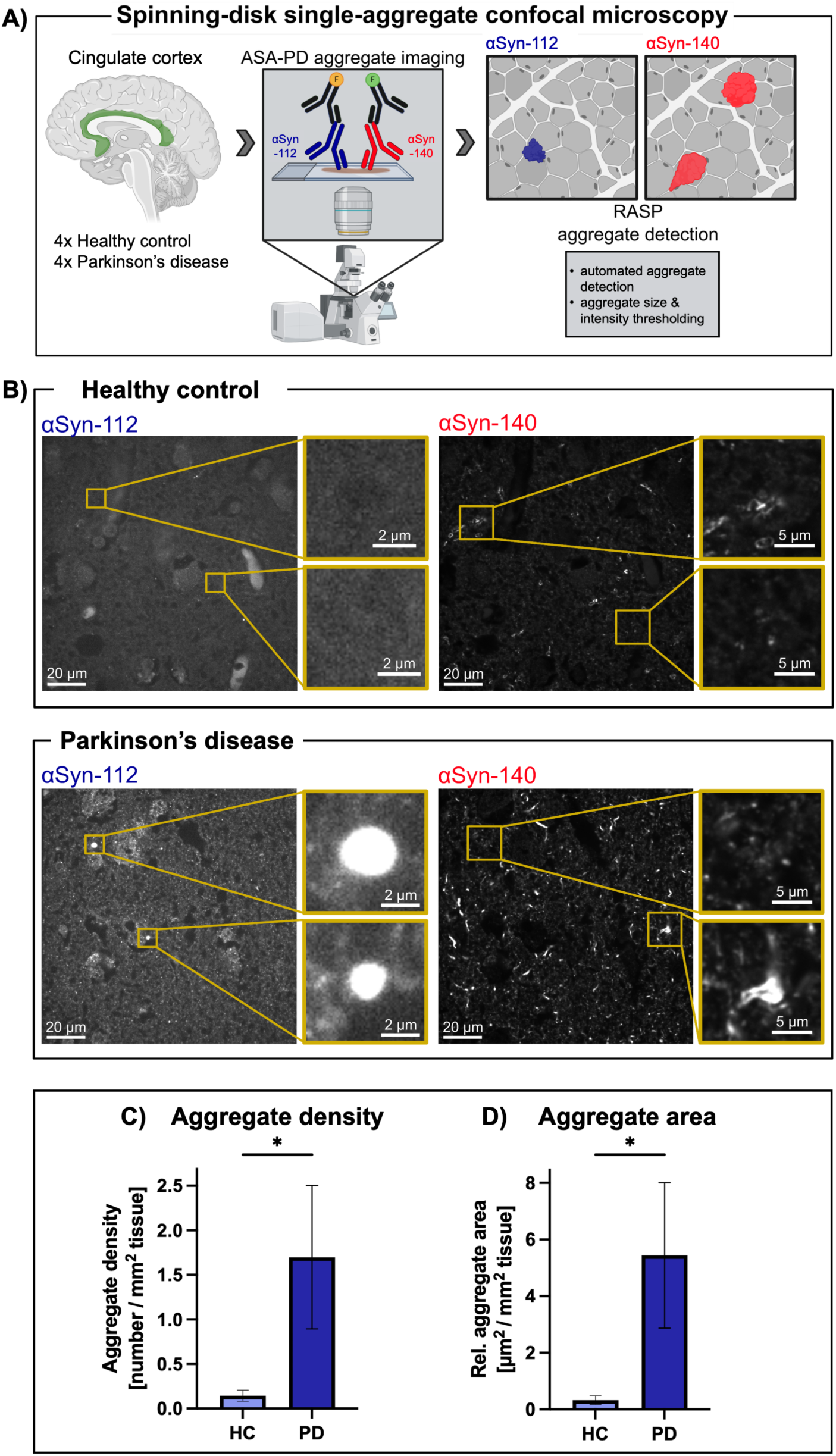
αSyn-112-immunoreactive aggregates are enriched in Parkinson’s disease brain tissue. **(A)** Conceptual workflow of αSyn detection in the cingulate cortex of healthy control and PD patients. **(B)** Representative microscopy images from healthy control (top) and PD (bottom) patients with detection of αSyn-112 (left) and αSyn-140 (right). Quantification of **(C)** the aggregate density and **(D)** the aggregate area of αSyn-112, normalised to the total tissue area analysed. Data are shown as mean ± SEM (n=4). Unpaired Mann-Whitney test. *p<0.05.

Using this approach, we found that both αSyn-112 and αSyn-140 immunoreactivity were increased in PD compared with control tissue (**Figure 6B**). The increase in αSyn-140 was expected from the established literature on αSyn deposition. For αSyn-112, immunoreactive aggregates were rare in control tissue, but markedly more frequent in PD (**Figure 6B**). A quantification confirmed this difference, as both the density and the relative area of αSyn-112-immunoreactive aggregates were significantly higher in PD than in controls (**Figure 6C,D**). In particular, the mean aggregate density was increased 12-fold (0.14 to 1.7 aggregates per mm^2^ of tissue area) and the relative aggregate area was increased 17-fold (0.32 to 5.44 µm^2^ per mm^2^ of tissue area), showing that these aggregates are both more numerous and larger in PD tissue. To our knowledge, this is the first IHC evidence that αSyn-112-immunoreactive deposits are enriched in PD brain. Moreover, in combination with the biophysical experiments conducted throughout this paper, these *in situ* results further corroborate our hypothesis that αSyn-112 may be a pathological factor in the development of synucleinopathies. Given that αSyn splice isoforms have not been considered a major pathogenic factor so far, our results open a new avenue of approaching and understanding the disease mechanism of PD.

## Discussion

In this study, we report that αSyn-112, an alternatively spliced αSyn isoform, exerts a significant influence on the aggregation of full-length αSyn-140. Although mounting evidence suggests that αSyn does not exist in the brain as a single species, but rather as a spectrum of proteoforms, alternatively spliced isoforms have received relatively little attention compared to point mutations and proteolytic truncations. Here, we describe how amounts of αSyn-112 as low as 1% are sufficient to significantly accelerate the aggregation of αSyn-140. Notably, this low amount lies within the physiological range of the αSyn-112 isoform, as long-read transcriptomics estimate αSyn-112 at up to 2.5% of total *SNCA* mRNA, representing the most abundant αSyn splice isoform after αSyn-140. Moreover, at the protein level, αSyn-112 has been detected in human brain and blood using top-down proteomics (*23, 24*). Furthermore, we provide evidence that this acceleration originates in enhanced heterotypic interactions between the two isoforms at the monomeric level, and that aggregated αSyn-112 is enriched in PD brain tissue relative to healthy controls. Taken together, these observations identify alternative splicing as a potent modulator of αSyn aggregation and motivate a closer examination of how the proteoform composition of αSyn shapes the initiation of pathology. To our knowledge, these results provide the first evidence that an alternative αSyn splice isoform can promote the overall aggregation of αSyn and, consequently, might be a key contributor to PD pathogenesis.

To elucidate the molecular mechanism of the co-aggregation between αSyn-140 and αSyn-112, we first performed a kinetic analysis and investigated the different microscopic steps that underlie the aggregation reaction. We found that the enhanced aggregation behaviour of αSyn-140+αSyn-112 mixtures *in vitro* is associated with an increase in primary nucleation, *i.e.* the molecular event by which protein monomers come together to form oligomers (*31*) (**Figures 1 and 2**). We thus sought to determine what types of oligomeric nuclei are initially formed. To probe these interactions, we employed MALDI-MS, a highly sensitive technique which allows for the detection of low-weight assemblies during the aggregation process. This approach revealed that αSyn-112 entirely formed hetero-oligomers with αSyn-140. Interestingly, no homo-oligomers of αSyn-112 were detected in this mixture, which is consistent with a preferential interaction between αSyn-112 with αSyn-140 monomers (**Figure 3**). Thus, using MALDI-MS, we provide direct evidence that two different splice isoforms of αSyn form hetero-oligomers, which in turn leads to an enhanced aggregation process.

We next performed NMR experiments to understand the molecular determinants of the enhanced monomer-monomer interaction between the two αSyn isoforms. We found that the deletion of the C-terminal exon 5 promotes the interaction between the NAC regions of αSyn-112 and αSyn-140 monomers. This is in agreement with literature showing that the hydrophobic NAC region drives αSyn aggregation (*9–11*). Moreover, the lack of the negatively charged exon 5 in the αSyn C-terminus may lead to reduced charge repulsion between monomers, which could explain why αSyn-112 monomers interact more strongly with αSyn-140 monomers compared to αSyn-140 monomers interacting with themselves. In addition, at least partially, αSyn-112 appears to engage with αSyn-140 in a head-to-tail orientation (**Figure 4**). This arrangement has previously been found in mixtures of αSyn with β-synuclein (βSyn), although we should note that this interaction was associated with an inhibitory effect on the aggregation with αSyn(*32, 33*). In line with the particular mode of interaction between αSyn-140 and αSyn-112 that we report here, hybrid αSyn-140+αSyn-112 fibrils exhibited a distinct secondary structure profile as per infrared spectroscopy (**Figure 5A,B**). This further corroborates the idea that the final morphology and structure of amyloid aggregates is highly dependent on the isoform composition, showcasing the effect that αSyn-112 can have on αSyn-140 in the context of disease-relevant protein aggregation. Integrating our findings across the various biophysical experiments shown in this study, the proposed model of co-aggregation and acceleration of aggregation is summarised in **Figure 7**.

**Figure 7.**
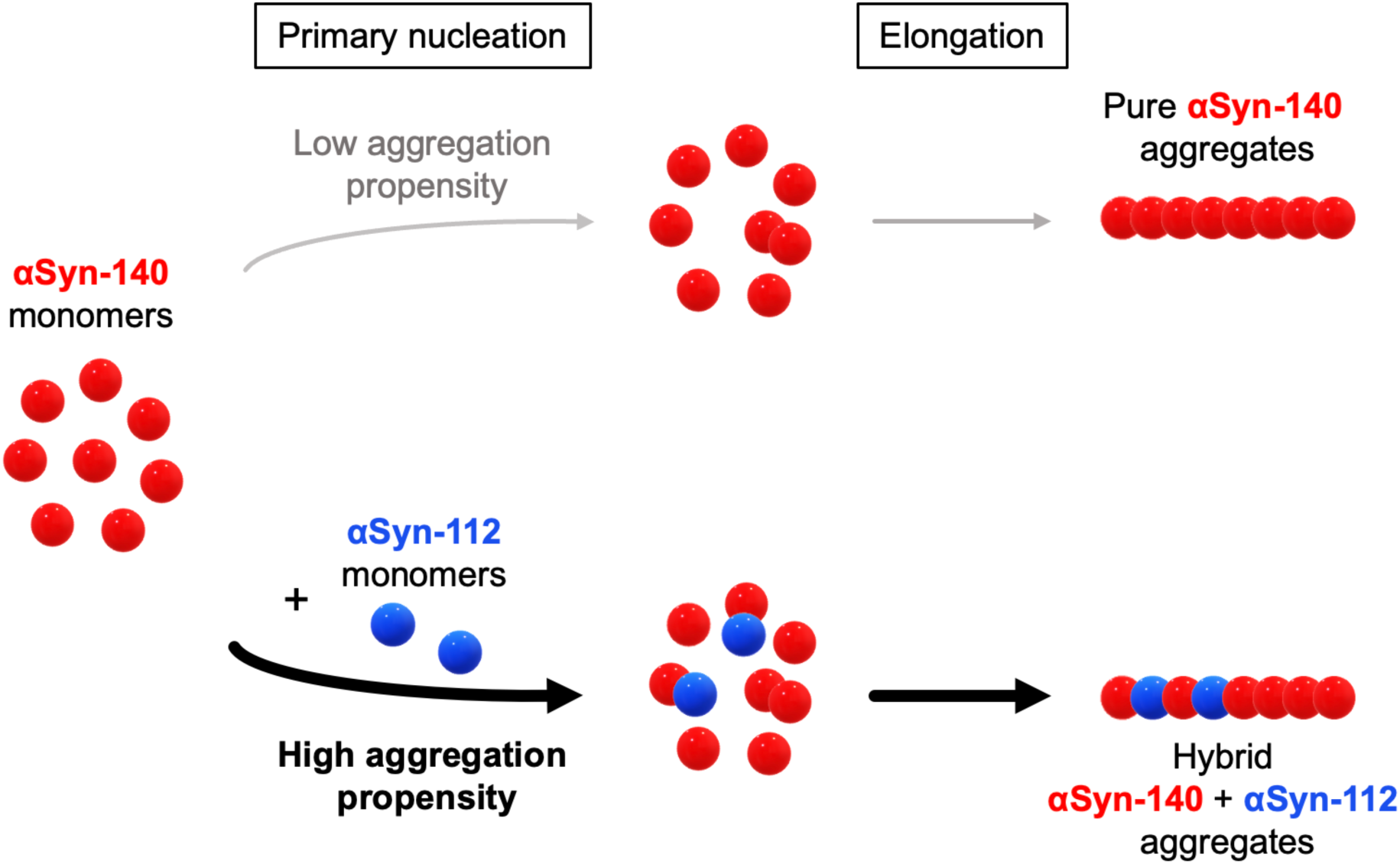
Proposed mechanism by which αSyn-112 promotes the heterotypic nucleation of αSyn-140 aggregation. The conversion of soluble αSyn into amyloid fibrils proceeds through a sequence of microscopic steps, beginning with primary nucleation, the assembly of monomers into the initial aggregates, which is slow for αSyn-140 alone (top). When present even in small amounts, αSyn-112 (blue) engages αSyn-140 (red), promoting the formation of heterotypic nuclei and raising the aggregation propensity of the mixture (bottom). This accelerated nucleation propagates through elongation into hybrid fibrils that incorporate both isoforms, so that a minor change in the proteoform composition of the monomer pool exerts a sizeable effect on the overall rate of αSyn aggregation.

While the previous parts of this study demonstrated the role of isoform interactions at the monomeric level, we furthermore explored the competency of the resulting fibrillar seeds in promoting the aggregation of the main αSyn-140 variant. In this regard, several studies have indicated that cross-seeding of αSyn-140 monomers with seeds of alternative αSyn variants leads to less efficient fibril growth due to structural incompatibility between the variant seeds and αSyn-140 monomers. Therefore, self-seeding, *i.e.* using seeds of the same species, has been found to be most effective (*13, 25, 34*). However, in this study, we went one step further and assessed the seeding potency of the aggregates resulting from co-aggregation (*i.e.* the seeds formed via the co-aggregation of 90% αSyn-140 + 10% αSyn-112). Interestingly, we found that the hybrid αSyn-140+αSyn-112 fibrils not only form faster than pure αSyn-140 fibrils but, strikingly, these hybrid fibrils were equally potent in further seeding αSyn-140 monomers (**Figure 5C-E**). This property bears important implications for the spread of αSyn aggregates in cellular systems, which has become an increasingly relevant topic in the field of αSyn research, given that the propagation of disease-associated aggregates through the brain may drive disease progression (*35, 36*). Integrating this in the context of our findings, it is possible that αSyn-112 leads to accelerated fibril formation with αSyn-140 and, when released, the resulting fibrils may induce αSyn aggregation in neighbouring cells.

Overall, our study forms part of a greater effort in recent years to investigate the behaviour and aggregation of αSyn proteoforms and their relationship with the development of synucleinopathies (*15, 19, 25, 37–39*). Notably, the principle that proteoform composition, and not merely the total level of an amyloidogenic protein, influences amyloid pathology is already well established for other neurodegenerative diseases. In Alzheimer’s disease, the more aggregation-prone Aβ42 is the primary component of amyloid plaques despite Aβ40 being the more abundant variant, and a shift in the Aβ42/Aβ40 ratio is regarded as a pathogenic feature promoting the development of AD (*40–43*). Likewise, six splice isoforms of tau are expressed in distinct patterns across human brain regions, cell types and developmental stages, and individual tauopathies are each dominated by a particular subset of these splice isoforms (*44, 45*). Our findings suggest that an analogous logic may apply to αSyn, meaning that changes in the expression pattern of *SNCA* transcripts can contribute to the development of synucleinopathies.

Previous co-aggregation studies involving αSyn variants predominantly focused on single point mutations or proteolytic truncations, *i.e.* the cleavage of the αSyn amino-acid chain. In particular, truncations of the αSyn C-terminus have been shown to enhance the aggregation of αSyn, which is in line with our results for αSyn-112 lacking the C-terminal portion encoded by *SNCA* exon 5 (*12, 13*). Decline in the proteostatic system constitutes one pathway through which the production of truncated proteoforms can be enhanced, although this phenomenon has been associated with advanced age (*17, 46, 47*). In contrast, alternative αSyn splice isoforms were found to be expressed in human brain tissue as early as the foetal stage (*5, 21*). Consequently, pathogenic splice isoforms may be generated at any point across the lifespan rather than predominantly in ageing. The possibility that splicing generates pathogenic isoforms of αSyn that directly contribute to disease, however, has largely been overlooked until now.

Finally, to investigate the relationship of αSyn-112 with disease, we explored the presence of αSyn-112 in tissue slices of PD patient brains and healthy controls using immunohistochemistry. To our knowledge, we demonstrate here for the first time that αSyn-112 exhibits significantly enhanced aggregate formation in PD patient brain tissue as compared to healthy controls (**Figure 6**). Moreover, our findings are in line with data from both the transcript and protein level, which showed increased αSyn-112 expression in brain lysate of patients afflicted by synucleinopathies (*37, 48–51*). Thus, by combining mechanistic insights *in vitro* with our observations *in situ*, our work provides an initial framework of evidence for how an alternative αSyn splice isoform could participate in human disease. We should note, of course, that as with every study, ours too presents a couple of limitations. Firstly, while long-read transcriptomic data indicate that αSyn-112 is the predominant exon-5-deficient splice isoform, the relative contribution of splice- and cleavage-derived species to the signal we observe will require further molecular identification of the deposited material, for example by mass spectrometry. Secondly, the tissue analysis was performed on a relatively small number of cases (n=4 per group) at a single, end-stage time point (Braak stage 6). It therefore establishes an association between αSyn-112 deposition and disease rather than a causal or temporal relationship, and larger, stage-resolved cohorts will be required to accurately determine whether αSyn-112 accumulation precedes or follows the bulk of αSyn-140 deposition and cellular changes in PD pathogenesis (*52, 53*).

Additionally, in support of our results, independent evidence also points towards a pathogenic role of αSyn-112. One study reported that αSyn-112, but not αSyn-140, leads to complement activation *in vitro* and might thus contribute to inflammation in PD (*54*). Furthermore, αSyn-112 exhibited enhanced binding to liposomes and oligomerisation on purified synaptic membranes *in vitro*, and caused severely impaired vesicle recycling and morphological changes at synapses *in vivo* (*55*). More broadly, the aggregation of αSyn is influenced not only by interactions among isoforms, as we describe here, but also by other amyloidogenic proteins. We have recently shown that the phase transitions of αSyn splice isoforms, including αSyn-112, can be modulated by Aβ peptides, with which αSyn is increasingly recognised to form co-pathologies in various neurodegenerative conditions (*56–58*). Together, these observations are in line with a general model of protein aggregation, which states that protein aggregation does not happen in isolation and is a complex process, for which contributions of alternative isoforms such as that of αSyn-112 must be taken into account.

Moreover, in order to develop effective pharmaceuticals targeting the aetiology of PD, it is instrumental to focus on the right steps in the disease pathway. To date, no drug that modifies the disease course is available for treating synucleinopathies, which can largely be attributed to the lack of understanding of the molecular events causing these diseases in the brain. Thus, the ultimate purpose of characterising the molecular mechanisms of αSyn aggregation is to learn how to control the disease process, and our findings carry several implications to do so. So far, efforts to inhibit αSyn aggregation have largely focused on the full-length protein, whereas our results show that the proteoform composition of αSyn contributes to its aggregation behaviour. Because αSyn-112, a low-abundance splice isoform, can substantially accelerate αSyn-140 aggregation through heterotypic interactions, an inhibitor optimised against αSyn-140 self-assembly may not equally suppress the hetero-oligomeric pathway described here. Our results therefore support designing inhibitors against mixed-proteoform aggregation, which may represent a promising new avenue in combatting synucleinopathies. Besides, this approach may be complemented by rational design techniques, which have been increasingly applied in recent years to design binders against a variety of proteins involved in neurodegenerative disorders (*59–62*).

Alternative approaches could aim at inhibiting the cellular production of αSyn-112 at the mRNA level, for example through antisense oligonucleotides directed at *SNCA* splicing (*19*). Moreover, selective enhancement of αSyn-112 clearance could reduce its availability for co-aggregation. Adding further complexity, amyloid proteins can interact with various cellular components and biomolecules, including other proteins, peptides, lipids, small metabolites and metal ions (*63–67*). Thus, future efforts should be directed at fully exploring the pathophysiology of αSyn-112, including causal and longitudinal studies, and at assessing the potential of αSyn-112 as a candidate biomarker for PD. More generally, we propose that synucleinopathies are associated with a heterogeneous population of αSyn proteoforms, and that resolving this heterogeneity will be crucial for understanding the molecular basis of this devastating class of disorders.

## Acknowledgements

A.R. acknowledges funding from the European Union’s Horizon 2020 research and innovation programme under the Marie Skłodowska-Curie grant agreement number 956977, and the TWIN2PIPSA programme under the grant agreement number 101079147. Z.T. acknowledges funding from the Ron Thomson Research Fellowship in Alzheimer’s Disease, Pembroke College, Cambridge. This research was funded in whole or in part by Aligning Science Across Parkinson’s (ASAP-000509) through the Michael J. Fox Foundation for Parkinson’s Research. The authors acknowledge the EPSRC Underpinning Multi-User Equipment Call (EP/P030467/1) for funding of the electron microscopy facility (Yusuf Hamied Department of Chemistry, University of Cambridge), and the MjölnIR National IR Infrastructure at MAX IV Laboratory (Lund University) supported by the MultiPark Infrastructure Grant. Ethical approval for experiments involving human tissue samples is provided by the Imperial College London Multiple Sclerosis and Parkinson’s Tissue Bank under ethics approval 23/WA/0273.

## Materials and Methods

### Recombinant expression and purification of αSyn isoforms

αSyn isoforms were expressed in BL21(DE3) Gold *Escherichia coli* bacteria (Agilent) using the pT7-7 plasmid (αSyn-140, αSyn-140-Q24C, αSyn-140-A90C) or pET29a(+) plasmid (αSyn-112, αSyn-112-Q24C, αSyn-112-A90C) (GenScript Biotech) and were purified based on previously described protocols (*13, 25, 68, 69*). Uniformly ^15^N-labelled αSyn-140 (N-terminally acetylated) was produced by co-transforming BL21(DE3) Gold *E. coli* with the pT7-7 plasmid and a plasmid carrying the components of the NatB complex (Addgene). The bacteria were grown in isotope-enriched M9 minimal media containing 1 g/L ^15^N ammonium chloride and 1 g ^15^N IsoGro (Sigma), and ^15^N-αSyn-140 was purified analogously to before.

### Aggregation assays

αSyn isoforms were diluted to the desired concentration in 50 mM Tris-HCl, pH 7.4, with 50 µM thioflavin T (ThT). The protein solutions were transferred onto Corning 96-well Half-Area Black with Clear Flat Bottom Polystyrene Non-Binding Surface Microplates containing a 3-mm borosilicate bead (Merck) per well and incubated under quiescent conditions on a FLUOStar Omega plate reader (BMG Labtech) at 37 °C. For seeded aggregation, αSyn-140 (100%) or αSyn-140 (90%) + αSyn-112 (10%) were pre-aggregated at a total protein concentration of 50 µM, with 50 µM ThT, for 6 days. The resulting fibrils were recovered from the plate, harvested by centrifugation (21100 *g*, 20 min, room temperature), and resuspended in 50 mM Tris-HCl, pH 7.4, with 50 µM ThT. Fibril concentrations in monomer equivalents were calculated assuming full monomer conversion. Fibrils were sonicated at 10 μM using a Sonopuls HD2070 microtip sonicator (Bandelin). The resulting fibril seeds were then added to αSyn monomers and aggregation assays were performed as described before.

### Kinetic analysis

Aggregation data were normalised with respect to ThT fluorescence, and half-times (t_1/2_) were calculated based on the timepoint when the normalised traces reached 0.5. All assays were performed with at least three repeats, and median traces were chosen for representation. The normalised data were fitted using the AmyloFit 2.0 platform to determine the microscopic rate constants.

### Transmission electron microscopy

3-mm 300-mesh copper grids (EM Resolutions Ltd.) were glow-discharged using a GloQube Plus (Quorum Technologies). Following aggregation, protein samples were recovered, 3 µL were spotted on the grids for 40 s and blotted using a Whatman filter paper. Subsequently, samples were stained with 2% (w/v) uranyl acetate for 40 s, blotted again and air-dried. Finally, TEM images were acquired using an FEI Talos F200X G2 transmission electron microscope (ThermoFisher Scientific).

### Production of MTSL-labelled recombinant αSyn

PRE experiments in this work required four types of spin-labelled αSyn molecules, each carrying a (1-oxyl-2,2,5,5-tetramethyl-Δ3-pyrroline-3-methyl) methanethiosulfonate (MTSL) spin label (Merck). To attach the MTSL to αSyn-140 and αSyn-112, we generated cysteine variants of the two isoforms. Two spin positions were alternatively used in this work, including residue 90 (through the point mutation A90C) and residue 24 (through the point mutation Q24C). ^14^N-MTSL-labelled cysteine variants were produced for both αSyn-140 and αSyn-112 (A90C or Q24C), rendering a total of 4 different constructs. The labelling was performed via the maleimide reaction using MTSL. These spin molecules were allowed to react with the thiol moiety of Cys90 or Cys24 using a well-established reaction protocol (*70*). The labelled proteins were then purified from the excess of free dye using a PD10 desalting column with a Sephadex G25 matrix (GE Healthcare) and concentrated (Amicon Ultra Centricons Millipore). The MTSL-labelled constructs were individually mixed with ^15^N-αSyn-140 to measure inter-molecular PRE effects probing protein-protein interactions.

### Paramagnetic relaxation enhancement (PRE) through transverse relaxation measurements

Intermolecular PRE was measured on ^15^N-αSyn samples at a concentration of 400 μM and incubated with each of the ^14^N-MTSL-labelled αSyn samples (400 μM). Relaxation enhancement in the NMR peaks of ^15^N-αSyn-140 indicated transient intermolecular contacts with the paramagnetic MTSL-labelled protein constructs. Relaxation was measured at 10 °C using a Bruker spectrometer operating at a ^1^H frequency of 700 MHz equipped with triple resonance HCN cryo-probe. Standard pulse sequences were used for T_2_ experiments(*71*), including the watergate sequence (*72*) to improve water suppression. Measurements were performed using a data matrix consisting of 2048 (t_2_, ^1^H) × 208 (t_1_, ^15^N) complex points and a recycle delay of 2.0 sec. CPMG delays were set as 16, 32, 64, 98, 128, 160, 192, 224, 288, 352, 416, 480, 544 and 608 ms, where spectra measured with the maximum delay showed less than 10% of signal intensity compared with those measured at 16 ms CPMG delay. Multiple data points were obtained for each CPMG delay. Data points were fitted with a single exponential function to provide T_2_ of individual amide N-H of the protein backbone. NMR spectra were acquired using Topspin 4.4.1 (Bruker) and processed with NMRpipe 10.9 (*73*) and Sparky 3.1 (*74*).

### Matrix-assisted laser desorption/ionisation mass spectrometry

50 µM αSyn-140, 50 µM αSyn-112, or 45 µM αSyn-140 + 5 µM αSyn-112 were aggregated for 20 h as described above. The aggregated sample was recovered and mixed with 2,5-dihydroxyacetophenone matrix (15.2 mg/mL 2,5-DHAP with 4.5 mg/mL diammonium hydrogen citrate (DAC)) and 2% trifluoroacetic acid (TFA) in a 1:1:1 ratio. 2 µL per sample were then spotted on polished steel plates and spectra were acquired on an ultrafleXtreme MALDI-TOF/TOF mass spectrometer (Bruker) in linear mode with positive polarity.

### Optical photothermal infrared spectroscopy

Optical photothermal infrared (OPTIR) spectra were collected using the mIRage Infrared Microscope (Photothermal Spectroscopy Corp.) as described previously (*75*). Specifically, OPTIR spectra were acquired in reflection mode with a 2 cm^−1^ spectral resolution, using a 40×, 0.78 NA Schwarzschild objective with an 8-mm working distance. The tunable four-stage quantum cascade laser (QCL) scanned from 1200 to 1800 cm^−1^, while the probe was a continuous-wave (CW) 532 nm laser with variable power. For sample preparation, a 1 μL drop of freshly prepared fibrillar suspension was placed on a glass substrate and dried under N_2_ flow. To achieve a sufficient signal-to-noise ratio, spectra were averaged over 5-10 scans. For data analysis, we used Quasar (*76*). First, the data were restricted to the Amide I or Amide III region, followed by rubber band baseline correction and min-max normalisation. The Average Spectra function was used to calculate the mean spectrum across all datasets, which served as the basis for defining the initial peak model. Peak fitting was subsequently performed for each individual spectrum using initial peak positions, amplitudes, and sigma values derived from the fitted average spectrum. Spectra were fitted by least-squares minimisation using user-defined composite Gaussian peak models. For each fitting parameter, minimum and maximum boundaries were defined as the initial value ± a specified delta. Initial peak positions were identified from the minima of the second-derivative spectra, and each component was modelled using a Gaussian function. The quality of the peak assignment was verified by comparing the second derivative of the fitted model with the experimental second-derivative spectrum, ensuring accurate identification of peak centres. Following peak fitting, the area of each fitted Gaussian peak was calculated and plotted for quantitative comparison.

### Tissue staining and spinning-disk confocal microscopy

Cingulate cortex (CC) tissue sections from four human post-mortem Parkinson’s disease cases (Braak Stage 6) and four healthy control cases were cut at 8 μm and prepared using the ASA-PD preparation protocol (*77*). Sections were blocked for 30 min at room temperature using BSA. Antigen retrieval was performed using formic acid, for ten minutes followed by heat-mediated epitope retrieval in Tris-EDTA, pH 9. The tissue was first stained for αSyn-112 overnight at 4 °C using the EPR25115-65 antibody (ab288156, abcam; 1:100 dilution) followed by goat-anti-rabbit secondary antibody conjugated to Alexa Fluor 568 (A11011, Thermo Fisher Scientific; 1:200 dilution) for 1 h at room temperature. While the primary antibody could theoretically also bind other exon 5-deficient αSyn splice isoforms, recent transcriptomics results indicate that αSyn-112 is by far the most predominant exon 5-deficient isoform in cells(*19*). Sections were then treated with Sudan Black (0.1% for 10 mins) before being mounted using VectaShield Antifade Mounting Media and sealed using CoverGrip. Subsequently, the tissue was stained for αSyn-140 using the syn211 antibody (ab80627, abcam; 1:100 dilution) followed by goat-anti-mouse secondary antibody conjugated to Alexa Fluor 488 (A11001, Thermo Fisher Scientific; 1:200 dilution) under the same pretreatment conditions. Stained brain tissue sections were imaged on a spinning-disk confocal microscope (3i intelligent imaging) built on a Zeiss Axio Observer 7 Basic Marianas™. The microscope was equipped with a 200 mW, 488 nm laser (LuxX) and a 150 mW, 561 nm laser (OBIS). These lasers were housed in a beam combiner (3i intelligent imaging), which focused them into an optical fiber that sent the illumination light into a field flattener (Yokogawa-Uniformizer for CSUW). The excitation light was then passed into a spinning disk unit (50 μm-sized pinholes, Yokogawa CSU-W1 T2 Single Molecule Spinning Disk Confocal, SoRa Dual Microlens Disk) and then the microscope body (Zeiss Axio Observer 7 Basic Marianas Microscope with Definite Focus 3) using a dichroic mirror (FF01-440/521/607/700, Semrock). The fluorescence was filtered using either a FF01-525/45-25-STR filter (Semrock) in the case of 488 nm excitation or a FF02-617/73-25-STR filter (Semrock) in the case of 561 nm excitation. The fluorescence was then focused onto one of two sCMOS cameras (Prime 95B, Teledyne Photometrics). The objective lens was a Zeiss oil immersion objective (Alpha Plan-Apochromat 100×/1.46 NA Oil TIRF Objective, M27). The microscope was controlled using a PC (Dell-Acquisition Workstation 310R) and SlideBook software produced by the manufacturer (3i intelligent imaging). For each tissue section, three x-y-z scans were acquired in random positions in the grey matter, and aggregate quantification of both αSyn-112 and αSyn-140 was achieved using the RASP pipeline (*30*).

## Notes

### Competing Interest Statement

The authors have declared no competing interest.

